# Plug and play: Is “directed endosymbiosis” of chloroplasts possible?

**DOI:** 10.1101/2021.12.03.471169

**Authors:** Karin Olszewski Shapiro

## Abstract

The origin of mammalian mitochondria and plant chloroplasts is thought to be endosymbiosis. Millennia ago, a bacterium related to typhus-causing bacteria may have been consumed by a proto-eukaryote and over time evolved into an organelle inside eukaryotic cells, known as a mitochondrion. The plant chloroplast is believed to have evolved in a similar fashion from cyanobacteria. This project attempted to use “directed endosymbiosis” (my term) to investigate if chloroplasts can be taken up by a land animal and continue to function. It has been shown previously that mouse fibroblasts could incorporate isolated chloroplasts when co-cultured. Photosynthetic bacteria containing chloroplasts have been successfully injected into zebrafish embryos, mammalian cells, and ischemic rodent hearts. The photosynthetic alga *Chlamydomonas reinhardtii* (*C. reinhardtii*) has also been injected into zebrafish embryos. However, to the best of my knowledge, injection of isolated chloroplasts into a land animal embryo has not been attempted before.

In four pilot experiments, solutions of chloroplasts in PBS were microinjected into *Drosophila melanogaster (D. melanogaster)* embryos to determine if the embryos would tolerate the foreign protein. Interestingly, results indicated that a portion of the *D. melanogaster* embryos appeared to tolerate the injections and survive to adulthood. To determine if chloroplasts had indeed been transferred, larvae were placed under fluorescent microscopy. Chlorophyll (serving as the reporter) was found to be present in several larvae and to decline in amount over time. To investigate if the chloroplasts still functioned, a radiotracer food intake assay was performed. It was hypothesized that if the chloroplasts were generating ATP (and possibly glucose), the larvae might need less food. Results indicated a decrease in intake, however this might have occurred for other reasons.

## Introduction

Humans have three energy systems: an aerobic process involving endosymbiotic mitochondria which provides us with the majority of our ATP supply; anaerobic glycolysis which breaks down glycogen into glucose when oxygen levels are low; and the anaerobic phosphocreatine system which uses muscle phosphocreatine to produce ATP. Plants have two endosymbiotic systems: chloroplasts which use light energy to convert atmospheric CO2 to glucose in the Calvin cycle; and mitochondria.

Mitochondria might be viewed as a “double-edged sword” for eukaryotes. On the one hand, these organelles provide us with 13 times more ATP than anaerobic respiration.^1^ On the other, the oxidative phosphorylation step of ATP production results in generation of cell-damaging reactive oxygen species (ROS). In plants, the chloroplast structure contains “thylakoid membranes” housing chlorophyll pigments,^2,3^ and “stroma,” fluid-filled regions containing NADP+. Chlorophyll absorbs photons, exciting electrons which then reduce NADP+ to NADPH in the stroma. Chlorophyll regains its electrons when water is photolysed, releasing gaseous oxygen.^4^ Photolysis releases protons (H^+^) which flow against their concentration gradient from the stroma to the thylakoid lumen. The enzyme ATP synthase then uses the energy from the gradient to generate ATP in the stroma. It is hypothesized that if ATP supply could be “augmented” in an animal by chloroplasts, food intake might decrease and hence less ROS may be generated by mitochondria. This might result in less ROS damage, leading to possible health benefits. It is interesting to speculate if longevity might also be affected. There appears to be an inverse relationship between food intake and life span. This relationship has been observed since the 1930s in multiple species.^5,6^

Support for this experiment may arise from Dr. Christina Agapakis’ 2011 Harvard master’s thesis in which the cyanobacterium *Synechococcus elongatus* (*S. elongatus)* was successfully microinjected into zebrafish embryos, with survival of both.^7^ In addition, *D. melanogaster* has been shown to already contain the endosymbionts *Spiroplasma* and *Wolbachia*.^8^ Further, an example of ATP augmentation in a marine animal already exists in nature. The sea slug *Elysia chlorotica* is reported to ingest chloroplasts from the alga *Vaucheria litorea* and derive nourishment from chloroplast photosynthesis for up to nine months.^9^ Health benefits from an endosymbiosis-like procedure have been discovered for cardiovascular disease. Stanford researchers Cohen et al. injected photosynthetic cyanobacteria *S. elongatus* into ischemic rodent hearts. Surprisingly, the results were a 25-fold increase in oxygenation vs. ischemic nadir.^10^ The goal of this experiment is to create a biomedical implant or patch containing chloroplasts, which might result in improvement of human health.

## Materials and Methods

### Chloroplast Isolation

For all trials, chloroplasts were isolated from spinach leaves using the Minute Chloroplast Isolation Kit (Product no. CP-011) from Invent Biotechnologies, Inc. (Plymouth, MN, USA). Isolation was performed at the Binninger lab at Florida Atlantic University. The kit contains filter cartridges with pore sizes designed to select for intact chloroplasts (>90% intact). 1×10^6^ to 1 × 10^7^ chloroplasts are pelletized by centrifugation and extraneous plant tissue remains in the cartridge. A homogenous sample solution was prepared by suspending the chloroplasts in PBS. Viability of the chloroplasts was tested using a 2,6-Dichlorophenolindophenol (DPIP) colorimetric assay. DPIP 0.1% solution was obtained from Carolina Biological Supply Company, Burlington, NC, USA (Product no. 746863). DPIP can act as a substitute electron acceptor for the chloroplast photosynthetic electron transport chain (ETC). Photosynthesis normally uses NADP+. DPIP is a blue solution that turns clear as it becomes reduced. A clear color result should indicate photosynthesis function. Color change of the DPIP treated solution was checked both visually and by spectrophotometer (Thermo Fisher Spectronic 20D+) transmittance reading (wavelength was set to 605nm.^11,12^)

### Fly Stock and Microinjection

For Trials 1-4, microinjection was performed by Rainbow Transgenic Flies, Inc. (Camarillo, CA, USA) (Rainbow Transgenic) using in-house stock male and female *D. melanogaster w*^*1118*^ which is a commonly used mutant strain (the mutation is in the *w* gene of the eye pigmentation pathway). At the time of injection, the flies were one-half to one hour old. Samples were injected in the germline (posterior) area.

In Trial 1, Rainbow Transgenic centrifuged a portion of the sample 1mL chloroplast/PBS solution using a Beckman Microfuge 16 centrifuge set at 5000rpm (equivalent to 1845g) for two minutes. No control was used as the purpose for this trial was to see if the injection was technically possible. Two groups of flies were injected as follows: (1) 134 embryos with non-spun chloroplast/1mL PBS solution; and (2) 130 embryos with spun chloroplast/1mL PBS supernatant. The sample was shipped overnight to Rainbow Transgenic on November 30, 2020. The company performed the injection on December 10 and overnighted the larvae to me on December 14, resulting in a 15-day time lag between chloroplast isolation and observation.

In Trials 2-4, no centrifugation was done. Instead, dilution was increased from 1mL to 2mL, 3mL and 4mL groups. PBS only controls were added. Sample solutions were shipped overnight in cold-pack boxes. Injected larvae were returned overnight the day of injection, resulting in a much improved 2-day time lag.

### ^32^P-Labeled Food Intake Assay

The ^32^P-labeled food intake assay was performed at the Ja lab at Scripps Research Institute. Larvae delivered on agar plates were transferred to standard stock food bottles and placed in a 25°C incubator to await the optimal developmental stage for the assay. There appeared to be good survival, ranking from control group (best), 3mL (next) and 2mL (least). There were clear developmental differences between the groups, following the same ranking order. Since the differences did not normalize enough for homogenous testing, non-pupariating larvae (located on the food area of the bottles, not the sides) were floated from the food with a 20% sucrose solution, then collected by pipette and rinsed with water. In this way, presence of pupae or wandering 3^rd^ instar larvae was avoided (larvae at this stage may reduce food intake). Larvae were transferred to ^32^P-labeled 2% yeast extract/5% sucrose food to perform the 6-hour assay. Individual larvae were then scintillation counted in 2.5mL fluid.

### Chlorophyll Fluorescence Microscopy

Fluorescence microscopy was performed at the McFarland lab at Florida Atlantic University, Harbor Branch Oceanographic Institute. Larvae were viewed using a Nikon Eclipse Ni-U microscope with an epifluorescence attachment, DS-Ri2 color camera, and installed filter cube (Chroma 49012 - ET - FITC/EGFP Longpass). The filter cube, when combined with the color camera, allows visualization of chlorophyll autofluorescence. Rainbow Transgenic provided the sample and control larvae taped to slides for ease of placement on the microscope stage.

## Results

Chloroplasts were isolated from baby spinach leaves using the Minute Chloroplast Isolation Kit, which selects for intact chloroplasts (>90% intact). Homogenous samples containing 1×10^6^ to 1 × 10^7^ chloroplasts were suspended in increasing dilutions of 1-3mL PBS (Fig 1).

**Fig 1.**
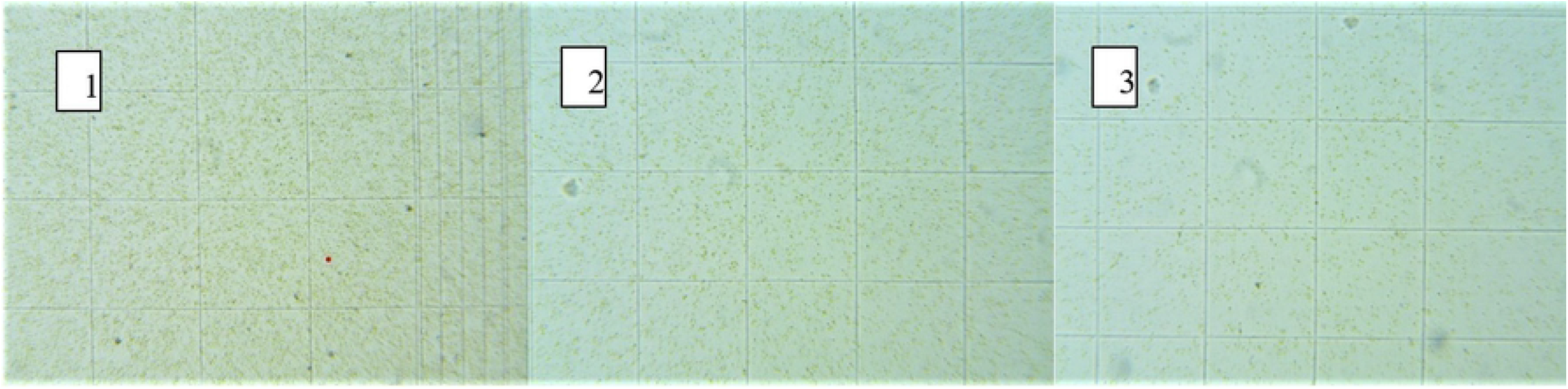
Chloroplasts Suspended in 1mL, 2mL and 3mL PBS. Image 1: Chloroplasts in 1mL PBS. Image 2: Chloroplasts in 2mL PBS. Image 3: Chloroplasts in 3mL PBS. Photos taken with an Amscope MU300 microscope camera on an Amscope M150C light microscope at 150x.

Viability of chloroplasts was tested using a DPIP colorimetric assay. DPIP can act as a substitute electron acceptor for the chloroplast ETC. DPIP is a blue solution that turns clear as it becomes reduced, therefore the clearer the result, the greater the photosynthetic function.^11,12^ Two solutions were prepared: (1) control: 600 uL ultrapure water + 6 drops of chloroplast/1mL PBS suspension; and (2) sample: control solution + 200uL 0.1% DPIP added. At time 11 minutes, the color of the sample cuvette had changed from dark to pale blue, indicating possible photosynthetic activity (Fig 2).

**Fig 2.**
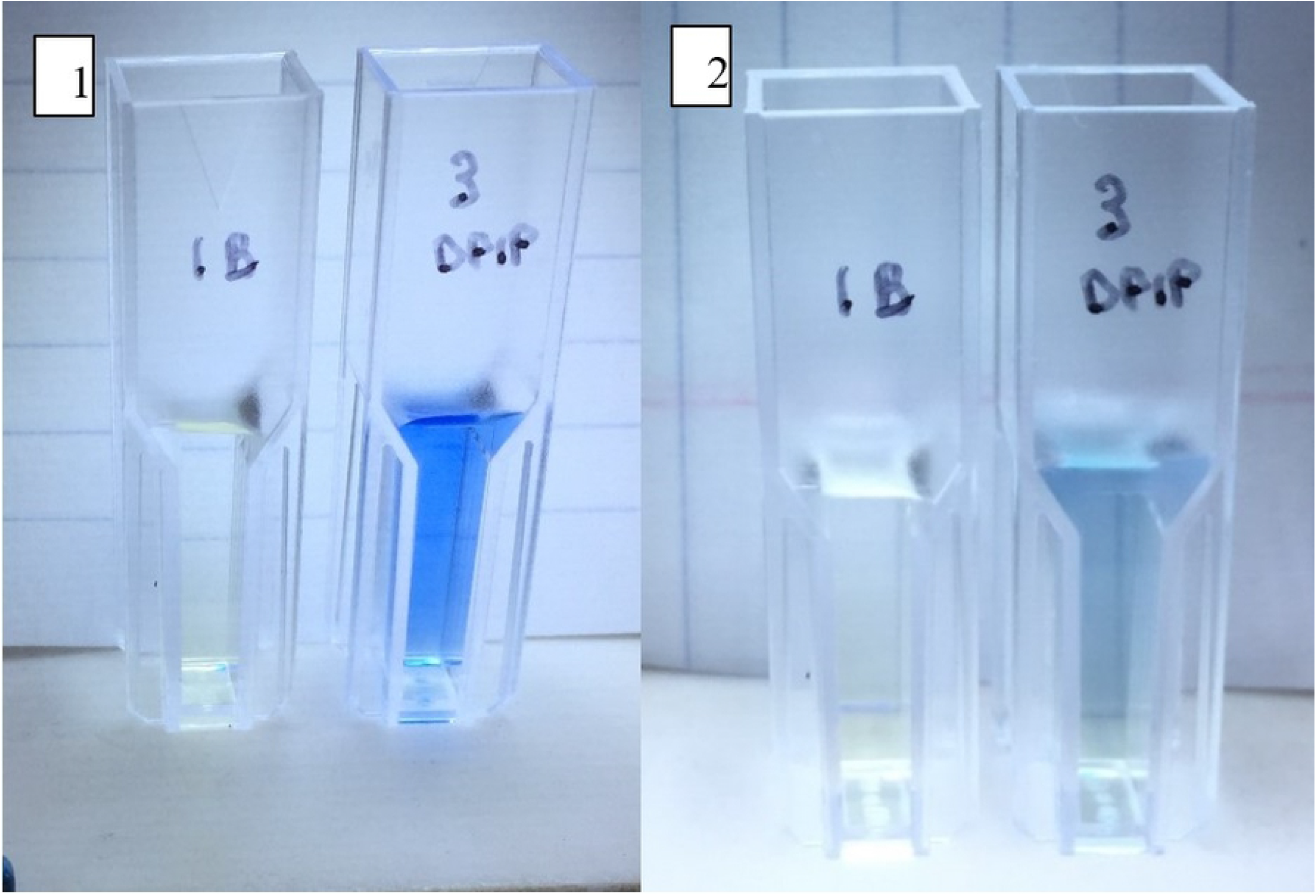
Visual Results of 0.1% DPIP Test on 1mL Chloroplast Solution. Image 1: Cuvette 1B (left) control solution - 600 uL ultrapure water + 6 drops of chloroplast/1mL PBS suspension at time 0. Cuvette 3DPIP (right) sample solution – control + 200uL 0.1% DPIP at time 0. Cuvette 1B is pale green. Cuvette 3DPIP is dark blue. Image 2: Cuvette 1B (left) control solution - 600 uL ultrapure water + 6 drops of chloroplast/1mL PBS suspension at time 11 minutes. Cuvette 3DPIP (right) sample solution – control + 200uL 0.1% DPIP at time 11 minutes. Cuvette 1B remained pale green. Cuvette 3DPIP appeared to change color from dark to pale blue which may indicate active photosynthesis.

Clarity of DPIP was also tested by spectrophotometry. An additional two solutions were prepared: (1) control: 4mL distilled water + 6 drops of chloroplast/1mL PBS suspension; and (2) sample: control solution + 200uL 0.1% DPIP added. At time 19 minutes, the spectrophotometer transmittance reading had changed from 40.2% to 74.0%, indicating possible photosynthetic activity (Fig 3).

**Fig 3.**
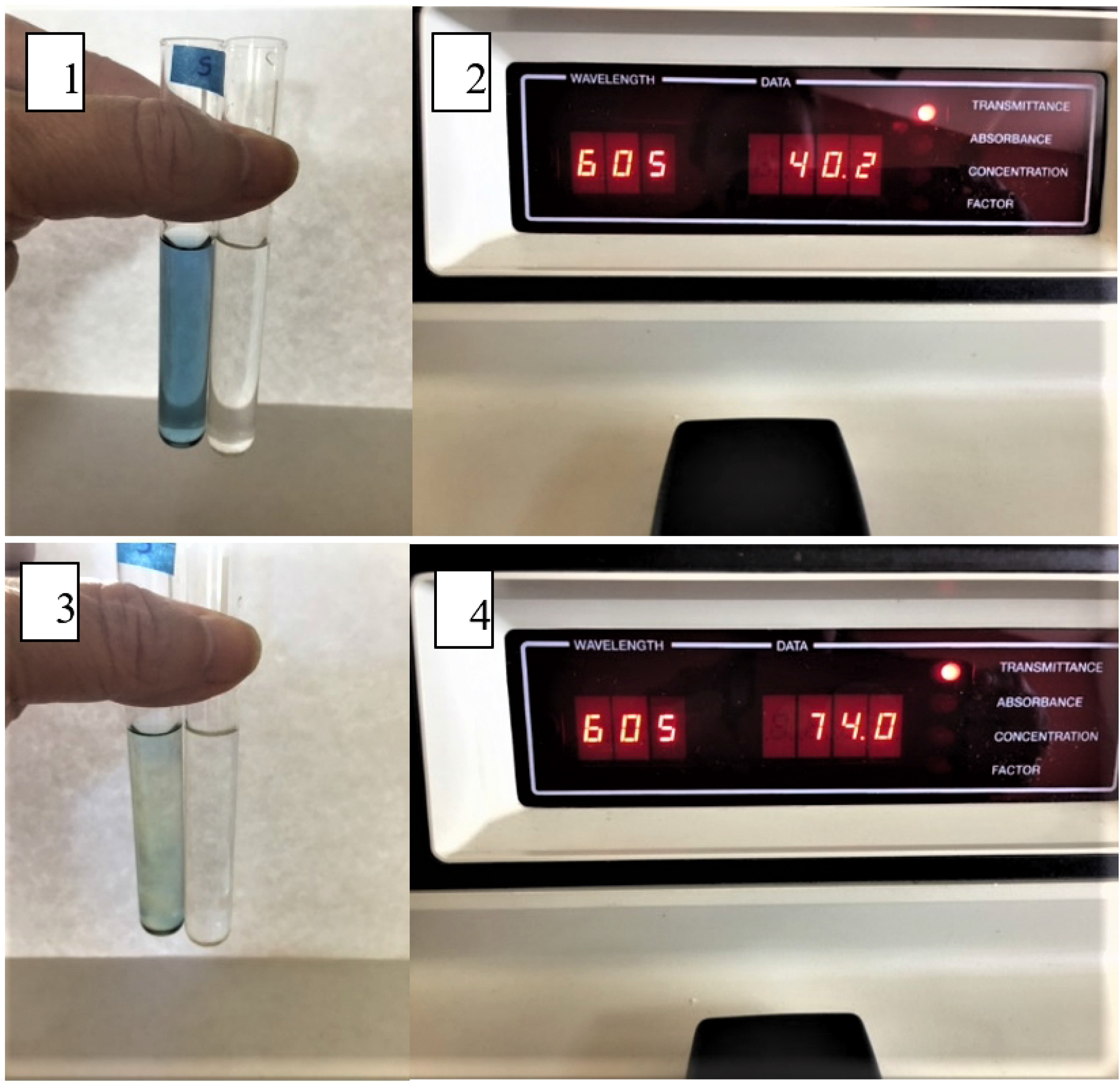
Spectrophotometer Results of 0.1% DPIP Test on 5mL Chloroplast Solution. Image 1: Vial S (left) sample solution – control + 200uL 0.1% DPIP at time 0. Vial C (right) control solution – 4mL distilled water + 6 drops of chloroplast/1mL PBS suspension at time 0. Image 2: Vial S spectrophotometer transmittance reading at time 0 – 40.2%. Image 3: Vial S (left) sample solution – control + 200uL 0.1% DPIP at time 19 minutes. Vial C (right) control solution - 4mL distilled water + 6 drops of chloroplast/1mL PBS suspension at time 19 minutes. Image 4: Vial S spectrophotometer transmittance reading at time 19.0 minutes – 74.0%. The control vial was used as the blank (transmittance set to 100%). The increase in light transmittance correlates with the solution becoming clearer as shown in Fig 2. This may indicate active photosynthesis.

### Trial 1

The purpose of Trial 1 was simply to see if microinjection of a chloroplast solution into *D. melanogaster* was technically possible. For this reason, a control solution was not included. A chloroplast/1mL PBS suspension was sent overnight to Rainbow Transgenic at room temperature. The company divided it into two samples – one centrifuged at 1845g for two minutes and the other not. It was found that the supernatant from the spun solution was markedly easier to inject than the non-spun one. 50 of 130 embryos survived the supernatant injection and 4 of 134 survived the non-spun injection. Out of the 50, approximately 30 ecloded (emerged from pupae). Out of the 4, 2 ecloded. Both groups survived for approximately two and a half weeks which falls below their average half-life (point of 50% survival) of approximately 45 days^13^ (Fig 4). This result appeared to indicate that chloroplasts can be injected into fly embryos without immediate lethality. However, without microscopy, it was not possible to know if the spun solution pelleted all the chloroplasts. Please see below.

**Fig 4.**
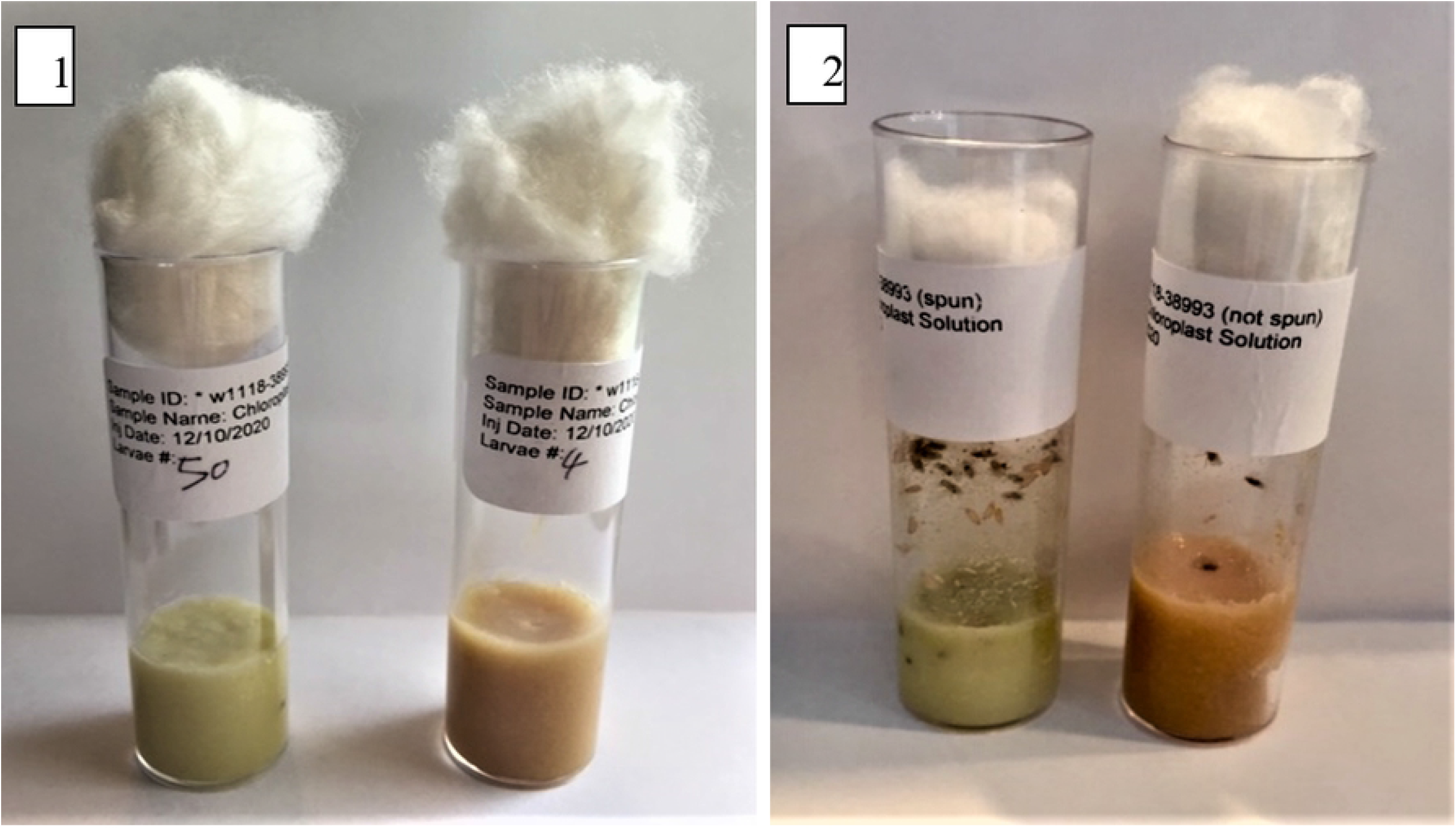
Photos of Vials Containing Injected *D. melanogaster*. Image 1: Vial on left contains supernatant injected embryos on Day 6 post-injection. 50 embryos out of 130 survived. Vial on right contains embryos injected with non-spun chloroplast 1mL solution on Day 6 post-injection. 4 embryos out of 134 survived. Image 2: Vial on left contains supernatant injected embryos on Day 18 post-injection. ∼30 embryos out of 50 ecloded. Vial on right contains embryos injected with non-spun chloroplast 1mL solution on Day 18. 2 embryos out of 4 ecloded.

### Trial 2 – Part 1

Building on the results of Trial 1, it was decided not to perform centrifugation. Instead, different dilutions were tested to observe which would work best. 2mL and 3mL suspensions, along with a PBS only control, were sent to Rainbow Transgenic overnight in a cold pack box. Rainbow Transgenic injected 250 embryos with the control, 315 with the 2mL dilution and 264 with the 3mL dilution. The company reported that the injections proceeded with increasing levels of difficulty, the control being the least and the 2mL dilution the most. Short video clips were taken of each injection group (Fig 5).

**Fig 5.**
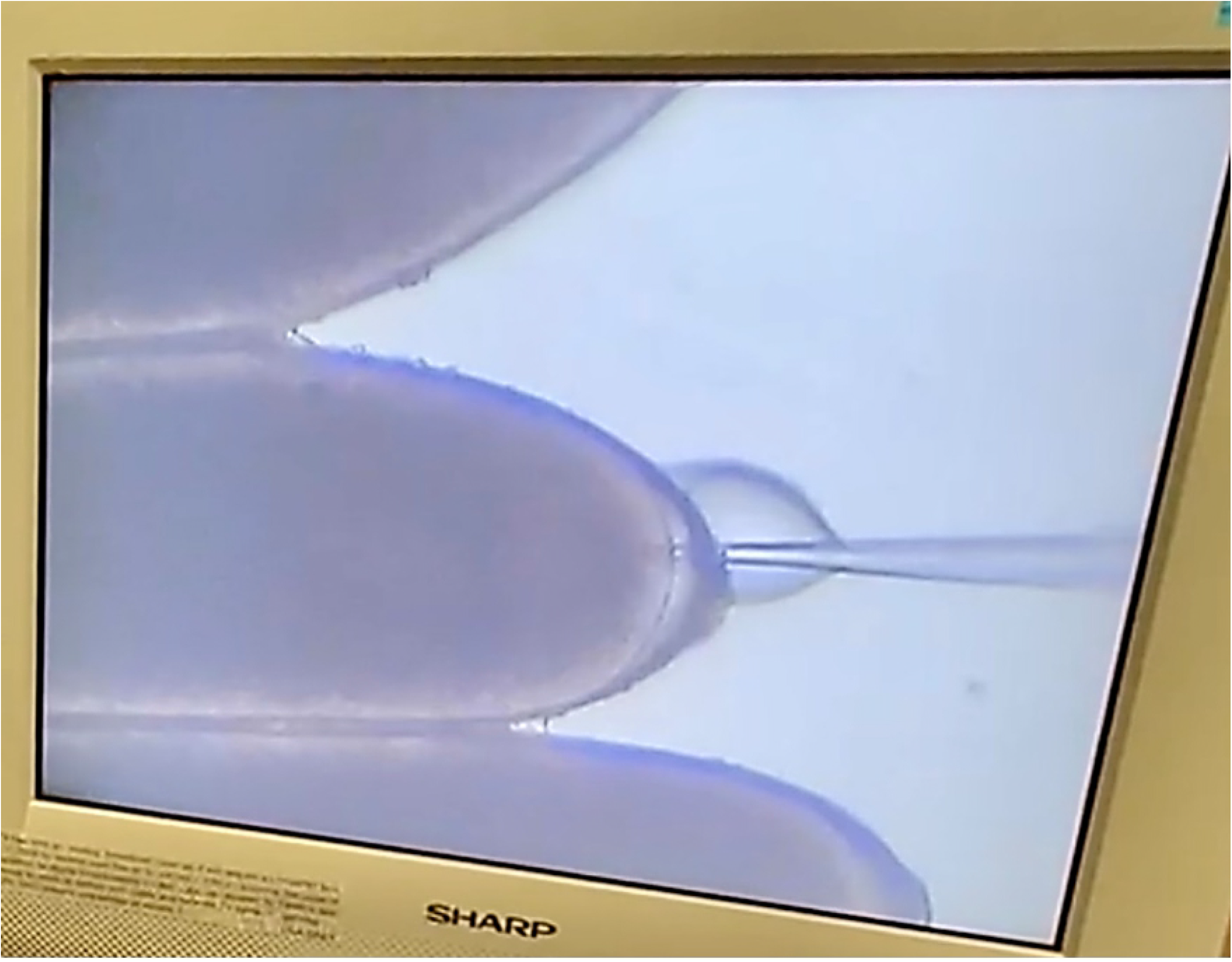
Screen Capture from Control Group Microinjection. Screen capture from video taken by Rainbow Transgenic on March 24, 2021.

Survival was compared for larvae in Trial 1 vs. Trial 2. The 1mL non-spun solution and the 2mL dilution had similar survival rates (3.0% and 3.8%, respectively). The survival rate for the 3mL dilution was ∽3.2 fold higher than for the 2mL dilution (Table 1). It was determined that 3mL appeared to be the best compromise to date between ease of injection and not over-diluting.

**Table 1.** Comparison of Larvae Survival - Trial 1 vs. Trial 2. The 1mL non-spun solution and the 2mL dilution had similar survival rates (3.0% and 3.8%, respectively). The survival rate for the 3mL dilution was ∼3.2 fold higher than for the 2mL dilution.

| **Trial 1 Larvae Sets**<br>**(Centrifuged sample solution)** | **Surviving**<br>**Larvae** | **Total Larvae** | **% Surviving Larvae** |
| --- | --- | --- | --- |
| Sample 1-Injected with supernatant from spun chloroplast/1mL PBS solution | 50 | 130 | 38.5 |
| Sample 2-Injected with non-spun chloroplast/1mL PBS solution | 4 | 134 | 3.0 |
| **Trial 2 Larvae Sets**<br>**(Diluted sample solutions)** | **Surviving**<br>**Larvae** | **Total Larvae** | **% Surviving Larvae** |
| Control 1-Injected with PBS only | 40 | 250 | 16.0 |
| Sample 1-Injected with chloroplast/2mL PBS solution | 12 | 315 | 3.8 |
| Sample 2-Injected with chloroplast/3mL PBS solution | 32 | 264 | 12.1 |

### Trial 2 – Part 2

Since survival improved over Trial 1, it was decided to test if the chloroplasts were possibly augmenting the embryos’ ATP supply. To indirectly determine this, food intake was measured. If food intake decreased, this might provide support for gain of energy from the chloroplasts.

Radio tracer studies on food intake for each group were performed as described in Materials and Methods above. Mean food intake was highest for the control (0.76mg), with the 3mL group next (0.49mg) and the 2mL group last (0.38mg). (Table 2 and Chart 1)

**Table 2.** Food Intake (mg) for Sample *D. melanogaster* Larvae. These larvae represent the portion that was floated from the food area with 20% sucrose. If max outliers are removed (highlighted in yellow), the averages change to 0.577613mg, 0.429605mg and 0.255171mg, respectively. (Source: Ja lab radio tracer study)

|  | Feeding (mg/larva) |  |  |
| --- | --- | --- | --- |
|  | Control | 3mL | 2mL |
|  | 0.045480 | 0.735730 | 0.120743 |
|  | 2.054650 | 0.718423 | 0.979631 |
|  | 1.058919 | 1.015049 | 0.153747 |
|  | 0.825482 | 1.354338 | 0.269258 |
|  | 0.114304 | 0.101424 | 0.596875 |
|  | 1.262573 | 0.042260 | 0.135233 |
|  | 0.147307 | 0.687030 |  |
|  | 0.589228 | 0.659661 |  |
|  |  | 0.765915 |  |
|  |  | 0.311518 |  |
|  |  | 0.217338 |  |
|  |  | 0.204861 |  |
|  |  | 0.085728 |  |
|  |  | 0.076068 |  |
|  |  | 0.768733 |  |
|  |  | 0.054335 |  |
| **Larvae no.** | 8 | 16 | 6 |
| **Mean** | 0.762243 | 0.487401 | 0.375914 |

**Chart 1.**
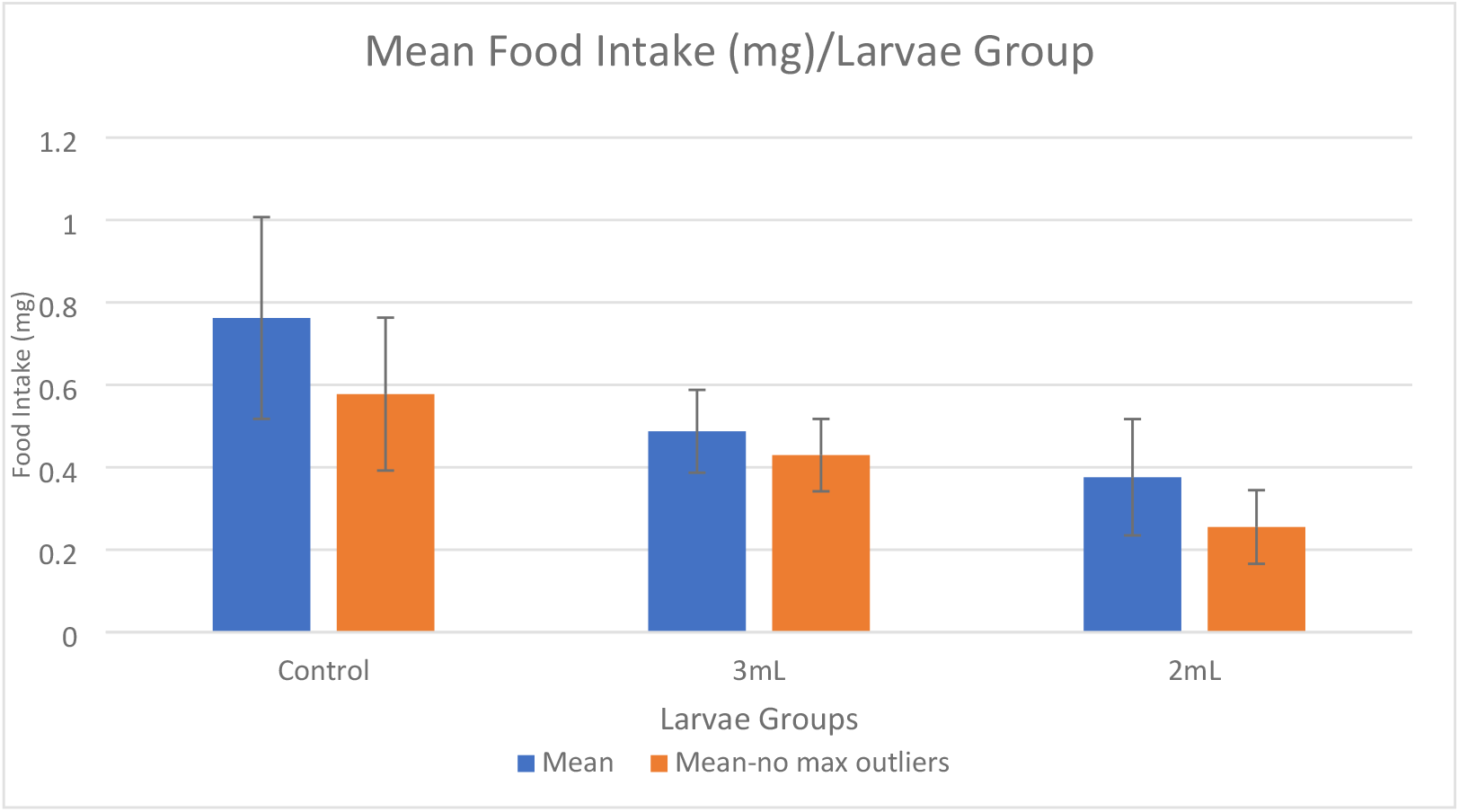
Chart 1. Mean Food Intake per Larvae Group – Standard Deviation. When max outliers are removed, standard deviation and error bars decrease. Mean food intake is highest for the control and successively decreases for the 3mL and 2mL groups. (Source: Adapted from Ja lab chart. I included additional column with max outliers deleted.)

As of April 12 (19 days post-injection), the non-food tested larvae appear to be thriving (Fig 6).

**Fig 6.**
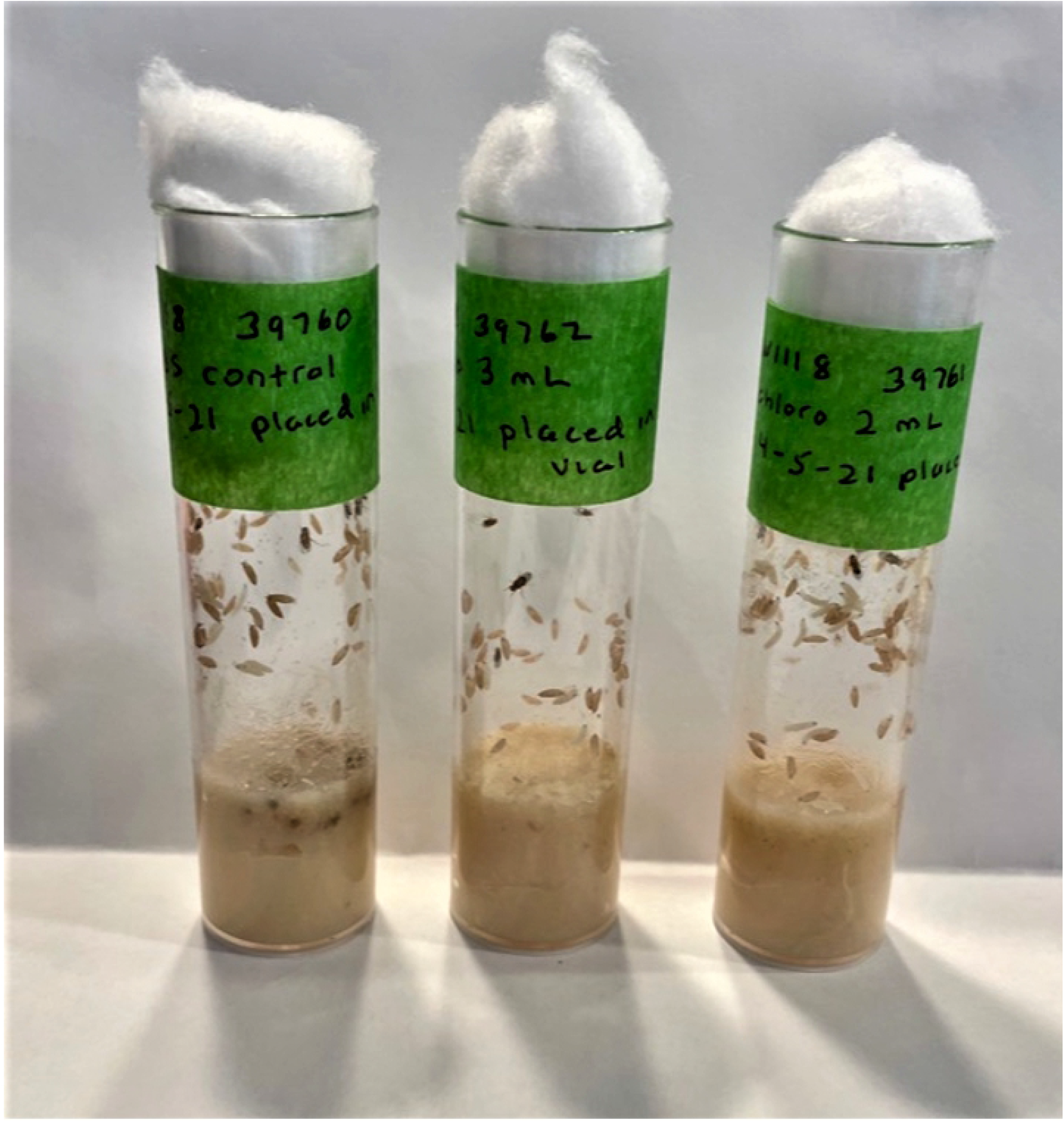
Non-Food Tested Larvae 19 Days Post-Injection. All three groups appear to be thriving.

### Trial 3

Fluorescence microscopy was performed to investigate if the microinjections successfully transferred chloroplasts into the embryos. Rainbow Transgenic provided four slides to which larvae had been taped as follows: (1) Control 1 – 100 non-injected larvae; (2) Control 2 – 270 larvae injected with PBS only; and (3) Sample - 160 larvae injected with chloroplast/3mL PBS solution divided among two slides. Rainbow Transgenic reported that it had been more difficult to inject the chloroplast/3mL PBS sample solution than in Trial 2. A possible cause might be slight variation in the amount of spinach leaf used for chloroplast extraction.

The slides were viewed under Lumencor Sola -generated blue light (450-490 nm) using a Nikon Eclipse Ni-U microscope.^14^ There appeared to be little interference from room ambient light. The expected result was that the larvae would appear green due to biological autofluorescence and any chlorophyll present would appear red. As in all living cells, larval autofluorescence is caused by the natural fluorescence of certain biological molecules such as pyridine nucleotides and flavins.^15^ Chlorophyll absorbs blue light which can be dissipated as heat, stored or used in other processes. The remaining blue light is emitted as longer wavelength red light.^16^

When viewed with the color camera, Control 1 (non-injected) appeared green with no visible red dots. Control 2 (PBS only) visualized green, with one red dot seen in one of the larvae. This may have been due to possible cross-contamination. The chloroplast/3mL PBS sample appeared green with a few scattered instances of red dots. Only one larva in this group showed presence of multiple red dots. (Fig 7) This result may have been caused by several factors. The chloroplasts may possibly have been degraded to some extent by the nascent immune system.^17,18^ Embryonic development during the approximate 2-day transit time transitions from syncytial to multicellular (cellularization).^18^ This might have interfered with the chloroplasts in a manner yet to be determined. Chloroplasts are no longer free-living organisms and may degrade in an unadapted host. In addition, prolonged exposure to light causes “photobleaching” of chlorophyll due to repeated cycles of excitation/emission. Fluorophores have a finite number of cycles before photon emission becomes disabled.^19^ The reported difficulty with injection may have contributed as well.

**Fig 7.**
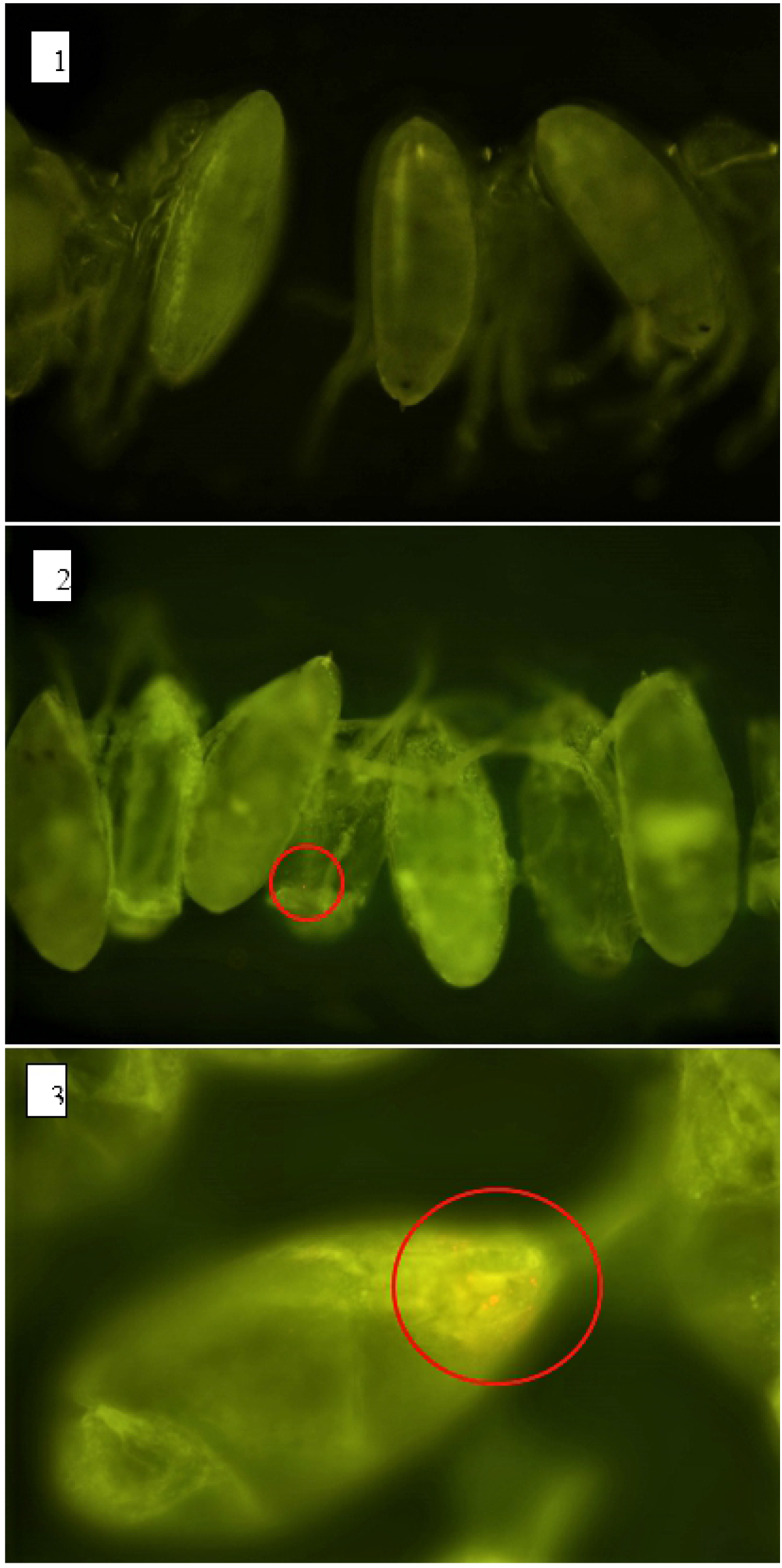
Images of *D. melanogaster* Sample and Controls Taken with Fluorescent Microscopy at a Resolution of 0.29 pixels um^-1^. Image 1: Control 1 larvae (non-injected) using 10x objective. Larvae appear green due to autofluorescence. No red dots indicating chlorophyll were observed. Image 2: Control 2 (PBS only) using 10x objective. Larvae appear green. One red dot observed near one larva (circled in red). Image 3: Sample (injected with chloroplast/3mL PBS solution) using 20x objective. Larvae appear green, with some scattered instances of red dots. One larva appeared to have multiple red dots (circled in red).

### Trial 4

The fluorescent microscopy procedure as described in Trial 3 was repeated, with the only difference being replacement of the 3mL dilution with a 4mL one to lessen potential difficulty of injection. Rainbow Transgenic provided four slides to which larvae had been taped as follows: (1) Control 1 – 235 non-injected larvae; (2) Control 2 - 260 larvae injected with PBS only; and (3) Sample - 250 larvae injected with chloroplast/4mL PBS solution divided among two slides. Rainbow Transgenic reported that injection went well.

When viewed with the color camera, Control 1 (non-injected) appeared green with visible red dots seen between two of the larvae. This may have been due to possible cross-contamination. Control 2 (PBS only) visualized green, with no visible red dots. The chloroplast/4mL PBS sample appeared green with instances of red dots on 27 larvae. Three larvae in this group showed presence of multiple red dots. (Fig 8)

**Fig 8.**
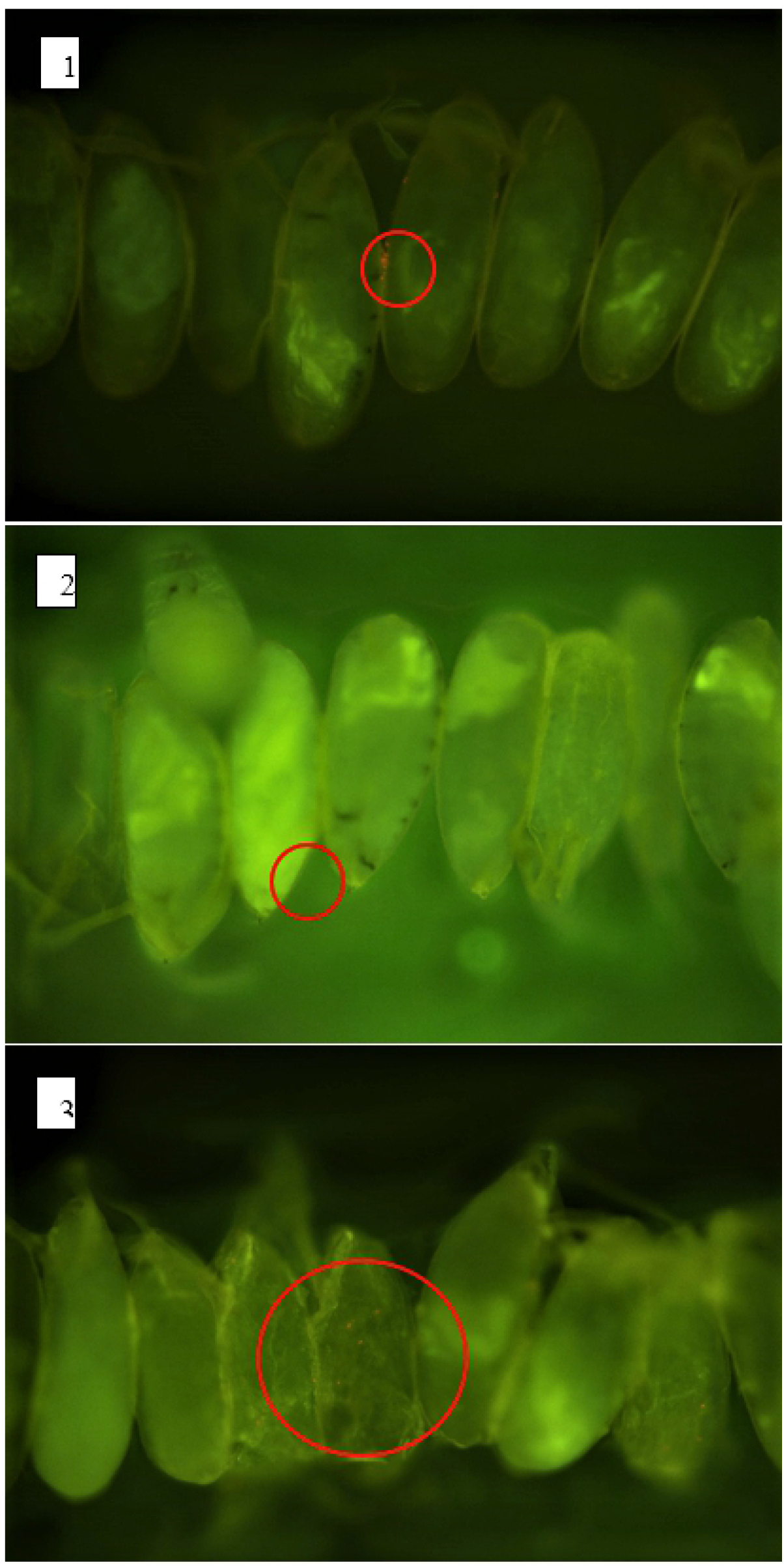
Images of *D. melanogaster* Sample and Controls Taken with Fluorescent Microscopy at a Resolution of 0.29 pixels um^-1^. Image 1: Control 1 larvae (non-injected) using 10x objective. Larvae appear green due to autofluorescence. One area of red dots (circled in red) was visible between two larvae indicating chlorophyll was observed. Image 2: Control 2 (PBS only) using 10x objective. Larvae appear green with no visible red dots. Image 3: Sample (injected with chloroplast/4mL PBS solution) using 10x objective. Larvae appear green. 27 larvae show visible red dots. Three of these larvae appeared to have multiple red dots (example circled in red).

The percentage of larvae showing presence of chlorophyll increased from 1.9% in Trial 3 to 10.8% in Trial 4 (Table 2). This ∼5.7-fold increase may be attributable to the greater dilution of the sample solution in Trial 4 reducing difficulty of injection and the decrease in time lag between chloroplast isolation and microscopy (two days vs. 4 days). The 50% reduction in time lag meant that the larvae may have been at an earlier state of cellularization, thereby lessening possible disruption of the chloroplasts. In addition, possible immune reaction would have been at an earlier stage. Finally, possible chloroplast degradation in an unadapted host may also have been at an earlier stage.

**Table 2.** Comparison of Fluorescent Microscopy Results - Trial 3 vs. Trial 4. The percentage of larvae showing presence of chlorophyll increased from 1.9% in Trial 3 to 10.8% in Trial 4.

| **Trial 3 Larvae Sets<sup>a</sup>** | **Larvae with Chlorophyll** | **Total Larvae** | **% Larvae with Chlorophyll** |
| --- | --- | --- | --- |
| Control 1-Uninjected | 0 | 100 | 0 |
| Control 2-Injected with PBS only | 1 | 270 | 0.37 |
| Sample-Injected with chloroplast/3mL PBS solution | 3 | 160 | 1.9 |
| **Trial 4 Larvae Sets<sup>b</sup>** | **Larvae with Chlorophyll** | **Total Larvae** | **% Larvae with Chlorophyll** |
| Control 1-Uninjected | 1 | 235 | 0.43 |
| Control 2-Injected with PBS only | 0 | 260 | 0 |
| Sample-Injected with chloroplast/4mL PBS solution | 27 | 250 | 10.8 |

<sup>a</sup>Time lag from chloroplast isolation to microscopy: 4 days.
<sup>b</sup>Time lag from chloroplast isolation to microscopy: 2 days.

## Discussion

### Embryo Survival

As shown in the Results section, survival rates increased in Trial 2 vs. Trial 1. Survival could not be tracked for Trials 3 and 4 due to embryo damage from fluorescent microscopy. However, it was observed microscopically (especially in Trial 4) that the embryos were moving and had structures forming inside. Since injecting a chloroplast solution into embryos is a new process both for the microinjection service and in general, a learning curve is involved. The average survival rate for standard forms of injection is ∼50%. As knowledge increases in terms of the best dilution and injection technique, it is expected that survival rates should increase.

An additional point is that mutant strain *w*^*1118*^ has been found to be stress intolerant under deficient or enriched nutrient conditions (stress simulation).^18^ One could extrapolate that despite this impairment, a portion of the larvae still survived. This might indicate that the injections may not have caused overwhelming stress even in such animals.

### Immune Reaction

These pilot studies had interesting results, particularly related to the apparent low level of immune reaction. One of the possible outcomes of Trial 1 could have been lethality of the test animals due to strong immune response to the foreign chloroplast proteins. The 3% and 12% survival rates in Trials 1 and 2, respectively, seem to belie this. However, a number of other issues could have caused this: accidental non-injection of some of the embryos, possible less developed immune response in larvae vs. adults,^17^ or the opposite – the larval immune system could have cleared the proteins. Detection of chlorophyll in Trials 3 and 4 indicate that a portion of the chloroplasts may have persisted for a certain amount of time. It is not possible to discern the exact role the immune system played without more sophisticated testing.

Three synthetic biology experiments mentioned previously showed apparent lack of an immune response to similar foreign proteins. These experiments involved injection of *S. elongatus* and *C. reinhardtii* into animal cells and *S. elongatus* into ischemic rodent hearts. Cyanobacteria are evolutionarily related to chloroplasts. Agapakis et al. found that *S. elongatus* did not seem immunogenic when injected into zebrafish embryos or mammalian CHO cells.^7,20^ Cohen et al. reported that *S. elongatus* did not arouse increase in CD8 T-cells, CD4 T-cells or CD19 B-cells in rodents.^10^ Alvarez et al. noted that *C. reinhardtii* also did not arouse immune response in zebrafish embryos as determined by observation of neutrophils under confocal microscopy.^21^ A fourth study by Nass found that isolated chloroplasts taken up by mouse fibroblasts remained in the cytoplasm, with no evidence of phagocytotic vesicles.^22^ The latter study is particularly interesting in light of the endosymbiotic theory. ^23^

### ATP Augmentation

The decrease in food intake between the control and the samples does not support or negate if ATP augmentation occurred. Although food intake might be hypothesized to decrease if this event happened, there was too much variability in the larval conditions to ascribe it to a particular cause. Possible reasons include growth and stage differences, amount of damage from injection, and relatively low number of test animals. Yet, a decrease *did* occur, therefore energy augmentation cannot be ruled out. Again, more sophisticated testing is needed to compare ATP levels in control vs. injected embryos at different time points.

## Conclusion

These pilot studies share a similar result with the four experiments mentioned above in that both isolated chloroplasts and cyanobacteria appear to survive initial introduction to a host. However, similar to the Nass experiment, the isolated chloroplasts decreased in number with time. It seems probable that the truncated genome of chloroplasts may be the cause. The majority of chloroplast genes have migrated to the host plant genome,^24^ which would likely affect viability inside an unadapted organism. It is planned to use synthetic biology techniques to construct a minimal genetic complement to extend viability. Migrated genes would be successively transformed into chloroplasts to determine the optimal assembly for increased longevity. The transformed chloroplasts would be injected into *D. melanogaster* embryos and ATP and glucose production measured *in vivo*. Test animals would be observed for biomarkers of immune response, oxygenation levels and life span/longevity. If results warrant, the next step would be to use a mammalian model. The eventual goal is to create an implant or patch for future biomedical use in humans.

## Acknowledgements

We thank the Binninger lab (Florida Atlantic University) for overall support, the Ja lab (Scripps Research Institute) for technical assistance with the radiotracer assays and the McFarland lab (Florida Atlantic University, Harbor Branch Oceanographic Institute) for assistance with fluorescence microscopy.

## References

1. Rich PR. The molecular machinery of Keilin’s respiratory chain. Biochemical Society Transactions. 2003;31(Pt 6):1095–1105. doi10.1042/BST0311095. PMID 14641005.

2. Dekker J, Boekema E. Supramolecular organization of thylakoid membrane proteins in green plants. Biochimica et Biophysica Acta (BBA) – Bioenergetics. 2005;1706(1–2):12–39. https://www.sciencedirect.com/science/article/pii/S0005272804002749

3. Von Wettstein D, Gough S, Kanangara GC. Chlorophyll biosynthesis. The Plant Cell. 1995; 7:1039–1057. https://www.ncbi.nlm.nih.gov/pmc/articles/PMC160907/

4. Royal Society of Chemistry. Photosynthesis. 2021. https://www.rsc.org/Education/Teachers/Resources/cfb/Photosynthesis.htm

5. McCay C, Crowell M. Prolonging the life span. The Scientific Monthly. 1934;39(5):405–414. www.jstor.org/stable/15813

6. McDonald R, Ramsey J. Honoring Clive McCay and 75 years of calorie restriction research. The Journal of Nutrition. 2010;140(7):1205–1210. https://doi.org/10.3945/jn.110.122804

7. Agapakis, C. Biological design principles for synthetic biology [M.Sc. Thesis]. Cambridge (MA): Harvard University; 2011. http://agapakis.com/thesis.pdf

8. Mateos M, Castrezana S, Nankivell B, Estes A, Markow T, Moran N. Heritable endosymbionts of Drosophila. Genetics. 2006;174(1):363–376. https://doi.org/10.1534/genetics.106.058818

9. Kenney D. Sea slug has taken genes from alga it eats, allowing it to photosynthesize like a plant. Phys.org. 2015. https://phys.org/news/2015-02-sea-slug-genes-algae-photosynthesize.html

10. Cohen J, Goldstone A, Paulsen M, Shudo Y, Steele A, Edwards B, et al. An innovative biologic system for photon-powered myocardium in the ischemic heart. Science Advances. 2017;3(6). e1603078. DOI: 10.1126/sciadv.1603078. https://advances.sciencemag.org/content/3/6/e1603078.full

11. Science Geek. Photosynthesis lab. https://www.sciencegeek.net/Biology/biopdfs/Lab_Photosynthesis.pdf

12. Carolina Biological Supply Company. Plant pigments and photosynthesis. https://www.ptbeach.com/cms/lib/NJ01000839/Centricity/Domain/113/ap%20biology%20Labs/apbiolab_04_plant_pigments.pdf

13. Ferreiro M, Perez C, Marchesano M, Ruiz S, Caputi A, Aguilera P, et al. Drosophila melanogaster white mutant w1118 undergo retinal degeneration. Frontiers in Neuroscience. 2018;11. https://www.frontiersin.org/articles/10.3389/fnins.2017.00732/full

14. Nikon Corporation. Nikon upright microscope Eclipse Ni-U instruction manual. https://www.manualslib.com/manual/837673/Nikon-Eclipse-Ni-U.html?page=10#manual

15. García-Plazaola JI, Fernández-Marín B, Duke SO, Hernández A, López-Arbeloa F, Becerril JM. Autofluorescence: Biological functions and technical applications. Plant Sci. 2015; 236:136–45. doi: 10.1016/j.plantsci.2015.03.010. PMID: 26025527. https://www.sciencedirect.com/science/article/pii/S0168945215000801?via%3Dihub

16. MiniPCR Bio. Chlorophyll lab: Green glows red. 2019. https://www.minipcr.com/wp-content/uploads/miniPCR-Chlorophyll-Glow_Lab_student_guide_v1.0_vF.pdf

17. Tan KL, Vlisidou I, Wood W. Ecdysone mediates the development of immunity in the Drosophila embryo. Current Biology: CB. 2014;24(10):1145–1152. https://doi.org/10.1016/j.cub.2014.03.062. https://www.ncbi.nlm.nih.gov/pmc/articles/PMC4030305/

18. Chaudhary R. Analysis of premature loss of the extraembryonic Amnioserosa in Drosophila morphogenetic mutants. Table 1.1. ResearchGate. 2021. https://www.researchgate.net/publication/266222693_Analysis_of_premature_loss_of_the_extraembryonic_Amnioserosa_in_Drosophila_morphogenetic_mutants

19. Davidson M. Photobleaching. Florida State University Optical Microscopy Primer. 2003. https://micro.magnet.fsu.edu/primer/java/fluorescence/photobleaching/

20. Agapakis CM, Niederholtmeyer H, Noche RR, Lieberman TD, Megason SG, Way JC, et al. Towards a synthetic chloroplast. PloS one. 2011;6(4). e18877. https://doi.org/10.1371/journal.pone.0018877. https://www.ncbi.nlm.nih.gov/pmc/articles/PMC3080389/

21. Alvarez M, Reynaert N, Chávez M, Aedo G, Araya F, Hopfner U, et al. Generation of viable plant-vertebrate chimeras. PLoS ONE. 2015;10(6). e0130295. https://doi.org/10.1371/journal.pone.0130295

22. Nass, M. Uptake of isolated chloroplasts by mammalian cells. Science. 1969;165(3898):1128–31. https://www.science.org/doi/10.1126/science.165.3898.1128

23. Sagan, L. On the origin of mitosing cells. Journal of Theoretical Biology. 1967;14(3):225–274. IN1-IN6. https://www.sciencedirect.com/science/article/pii/0022519367900793

24. Oliver JL, Marín A, Martínez-Zapater JM. Chloroplast genes transferred to the nuclear plant genome have adjusted to nuclear base composition and codon usage. Nucleic Acids Research. 1990;18(1):65–73. https://doi.org/10.1093/nar/18.1.65

